# A Bayesian observer model reveals a prior for natural daylights in hue perception

**DOI:** 10.1101/2023.06.06.543889

**Authors:** Yannan Su, Zhuanghua Shi, Thomas Wachtler

**Affiliations:** Faculty of Biology, Ludwig-Maximilians-Universität München, Planegg-Martinsried, Germany; Graduate School of Systemic Neurosciences, Ludwig-Maximilians-Universität München, Planegg-Martinsried, Germany; General and Experimental Psychology, Ludwig-Maximilians-Universität München, Munich, Germany; Bernstein Center for Computational Neuroscience Munich, Planegg-Martinsried, Germany

## Abstract

Incorporating statistical characteristics of stimuli in perceptual processing can be highly beneficial for reliable estimation from noisy sensory measurements but may generate perceptual bias. According to Bayesian inference, perceptual biases arise from integrating internal priors with noisy sensory inputs. We used a Bayesian observer model to derive biases and priors in hue perception based on discrimination data for hue ensembles with varying levels of chromatic noise. For isoluminant stimuli with hue defined by azimuth angle in cone-opponent color space, discrimination thresholds showed a bimodal pattern, with lowest thresholds near a non-cardinal blue-yellow axis that aligns closely with the variation of natural daylights. Perceptual biases showed zero crossings around this axis, indicating repulsion away from yellow and attraction towards blue. The biases could be explained by the Bayesian observer model through a non-uniform prior with a preference for blue. Our results suggest that visual processing exploits knowledge of the distribution of colors in natural environments for hue perception.

## 1 Introduction

The dynamical and statistical nature of the sensory environment poses challenges for sensory processing and perception. Sensory responses to the same stimulus can di↵er, and di↵erent stimuli can cause similar sensory stimulation. However, the natural sensory world is not entirely random but exhibits regularities, and exploiting this predictability can help an organism in taking useful decisions and efficient actions. Achieving this, however, requires that the knowledge about the sensory environment is incorporated in sensory processing.

That sensory processing might utilize knowledge about the regularities of the world can be traced back to von Helmholtz’s idea of ’unconscious inference’ (von Helmholtz, 1867). In recent decades, the development of the Bayesian inference framework suggests that incorporating prior knowledge can significantly enhance the reliability of perceptual estimation, especially when the input signals are corrupted by noise (Knill and Pouget, 2004; Shi et al., 2013). The Bayesian inference framework has successfully accounted for perceptual performance in object perception (Kersten et al., 2004), multi-sensory integration (Ernst and Banks, 2002), sensorimotor learning (Körding and Wolpert, 2004), visual speed perception (Stocker and Simoncelli, 2006), visual orientation perception (Girshick et al., 2011), and time perception (Shi and Burr, 2016; Glasauer and Shi, 2022).

In the case of orientation perception, Bayesian approaches have revealed that human observers’ perceptual judgments are systematically biased towards cardinal orientations (Girshick et al., 2011). The corresponding prior matched the non-uniform distribution of orientation in natural scenes, where orientations near the cardinals have a higher incidence than oblique orientations (Girshick et al., 2011). Non-uniformities in the statistics of natural sensory signals exist also in the domain of color. For example, distributions of cone-opponent signals in natural scenes show a correlation between S-(L+M) and (L-M) coordinates (Webster and Mollon, 1997; Wachtler et al., 2001; Nascimento et al., 2002; Webster et al., 2007), indicating a dominance of contrasts along an oblique color space axis that corresponds to the variation of natural daylights. Analyzing human color perception in a Bayesian framework may provide insights into how human color perception is adapted to such non-uniformities in the chromatic properties of the natural environment.

In the Bayesian framework, an observer’s statistical inference is influenced by two key components: the likelihood, which is the probability of sensory measurements given a stimulus, and the prior, which reflects the observer’s prior knowledge about stimulus probabilities (Knill and Pouget, 2004). Optimal integration of the two components results in a posterior density function that represents the probability of the stimulus given the measurements.

A common difficulty encountered in the Bayesian inference framework lies in measuring likelihood and prior, which are not directly accessible. Stocker and Simoncelli (2006) proposed a method to recover likelihood and prior from psychophysical measurements of perceptual bias and variability. Specifically, these measurements were obtained from discrimination between stimuli with different noise levels, corresponding to different widths of the likelihood. According to the Bayesian inference framework, this results in di↵erent posterior distributions. When the likelihood does not align with the prior, the larger the noise, the larger the shift in the posterior induced by the prior (Fig. 1). Girshick et al. (2011) used this approach to investigate orientation perception and revealed a prior that peaked at the cardinal orientations, suggesting that visual perception may involve prior information regarding the regularities of natural scene structures.

**Figure 1:**
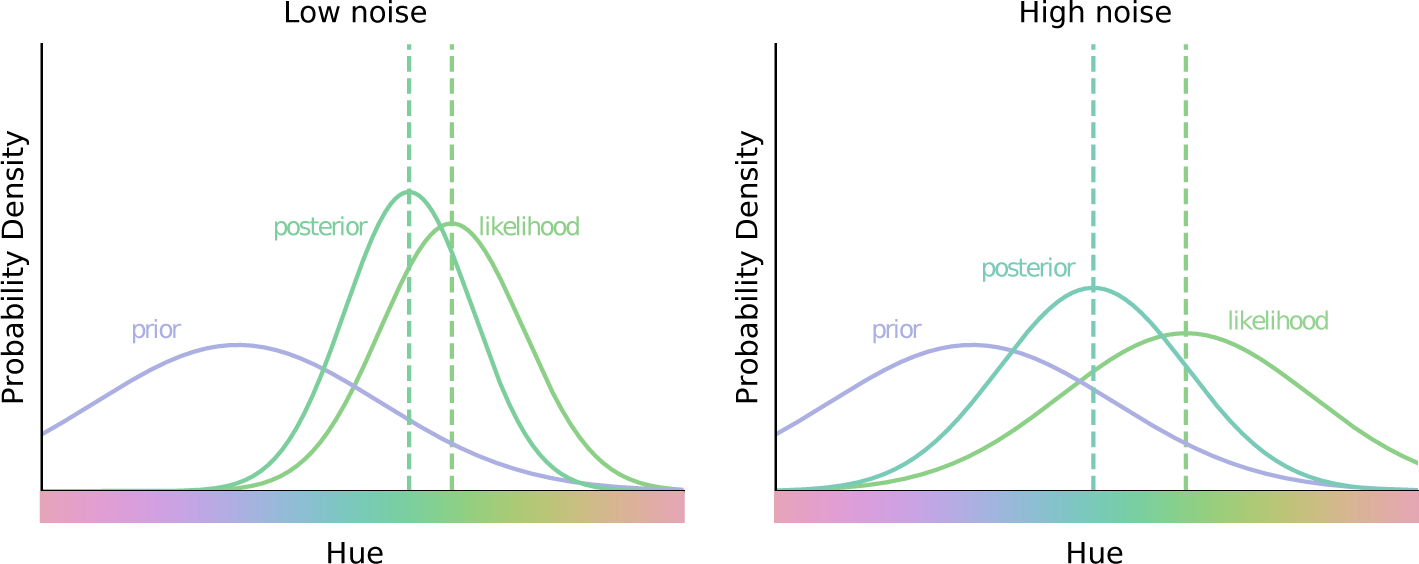
Illustration of Bayesian inference in hue perception (after Stocker and Simoncelli, 2006). The same prior integrates with the likelihoods for two stimuli with di↵erent noise levels. Left: A stimulus with a low level of noise results in a narrow likelihood and thus a small shift of the posterior. Right: A stimulus with a high level of noise results in a wide likelihood and thus a large shift of the posterior.

To investigate how prior knowledge is integrated into color processing, we closely followed the approach by Stocker and Simoncelli (2006) and Girshick et al. (2011), applying the Bayesian framework in the domain of hue perception. We measured perceptual variability and biases of observers in the discrimination of noisy hue ensembles and used a Bayesian model to recover their priors.

## 2 Methods

### 2.1 Participants

Six observers, (two males and four females), ranging in age from 23 to 56 years, participated in the experiments. All observers had normal or corrected-to-normal vision. Participants signed informed consent prior to the experiment and received a compensation of 10 Euros per hour. The study was conducted in accordance with the Declaration of Helsinki.

### 2.2 Stimuli

Stimuli were presented on a ViewPixx Lite 2000A display, calibrated by a PhotoResearch (Chatsworth, CA) PR-655 spectroradiometer, and controlled by a Radeon Pro WX 5100 graph-ics card. The screen resolution was set to 1920 *⇥* 1200 pixels at a refresh rate of 120 Hz.

Chromaticities of the stimuli were defined in a cone-opponent color space similar to the one used by Derrington et al. (1984), but with the y-axis representing increasing S-cone stimulation (MacLeod and Boynton, 1979). In this color space, the distance from the center corresponds to chromatic contrast, the azimuth angle corresponds to hue, and the orthogonal third axis corresponds to luminance contrast. Cone contrasts were defined with respect to a neutral gray (106.7 cd/*m*^2^, CIE [x, y] = [0.307, 0.314]), which was used as the display background on which stimuli were presented. The Scone contrast axis was scaled by a factor of 2.6, yielding approximately equally salient stimuli for all hues at a given cone contrast (Teufel and Wehrhahn, 2000). Individual perceptual isoluminance with respect to the reference gray was determined for 16 stimuli of different hues using heterochromatic flicker photometry (Kaiser and Boynton, 1996). From these data, an isoluminant plane in cone-opponent color space was calculated individually for each subject (Teufel and Wehrhahn, 2000). All stimuli used were isoluminant and had a fixed cone contrast of c=0.12 with respect to the neutral gray background. Thus, stimuli varied only in azimuth angle, corresponding to hue (Fig. 2a).

**Figure 2:**
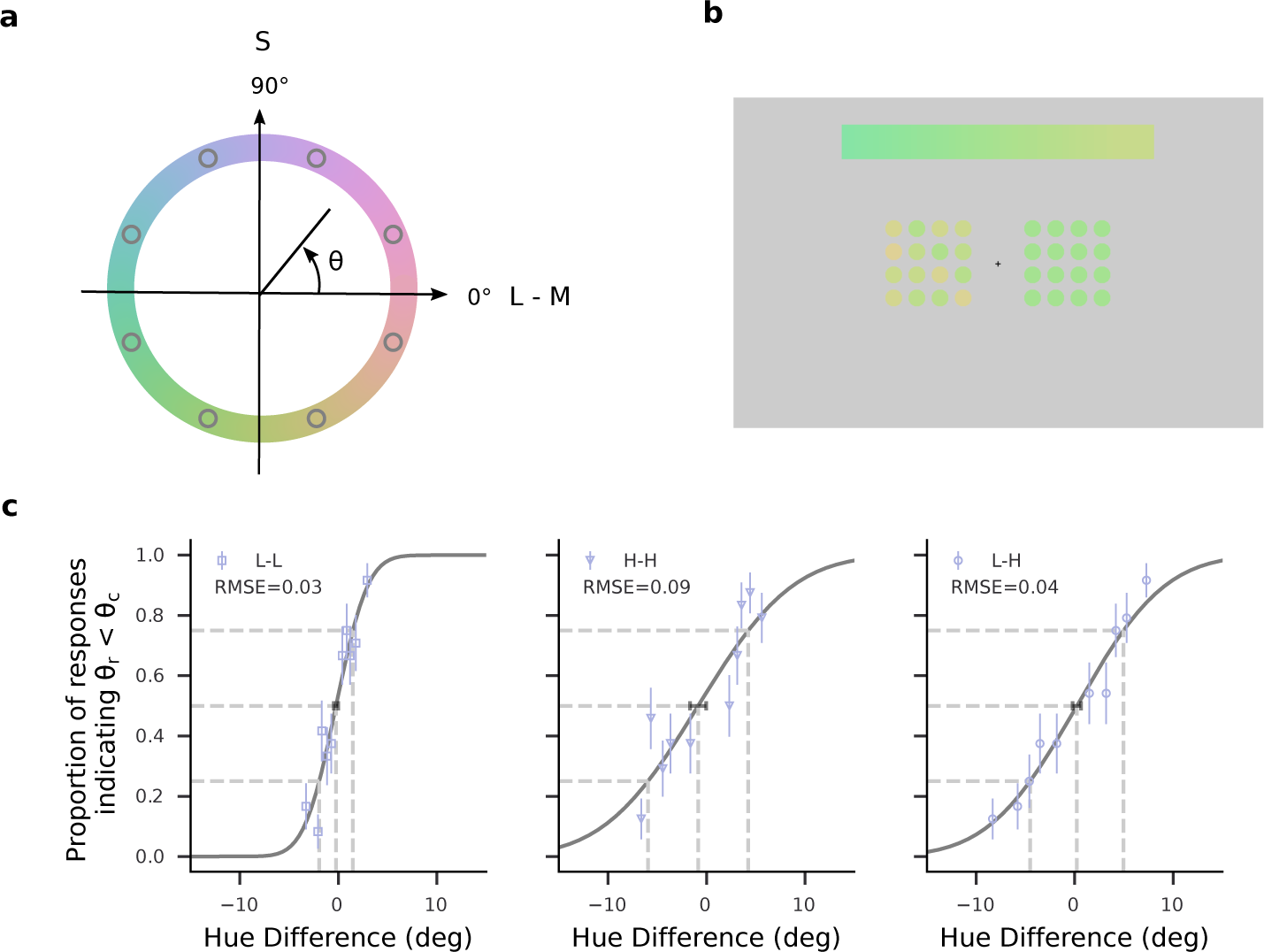
Stimuli and psychometric functions. (a) Stimulus chromaticities. All stimuli were defined in the isoluminant plane with fixed chromatic contrast and thus varied only in azimuth angle, corresponding to hue (colored circle). Eight hues with equidistant azimuth angles *θ* (gray open dots) were defined as reference hues. (b) Example of a stimulus display. Participants were asked to compare two arrays of color patches, presented on the left and right side of fixation, and to indicate whether their respective average hues matched the direction of hue changes that was indicated by the color gradient bar at the top of the display. The figure shows the cross-noise condition with a high-noise array on the left and a low-noise array on the right. Note that hue di↵erences have been exaggerated in the figure for illustrative purposes. (c) Examples of psychometric functions for the three noise conditions. Proportions of responses indicating the subject perceived the hue angle of the comparison stimulus (*θ_c_*) as larger than the hue angle of the reference stimulus (*θ_r_*) are plotted as a function of the di↵erence between the hue angles *θ_c_* and *θ_r_*. Data are from a single subject’s responses with *θ_r_* = 112.5° for the three noise conditions L-L (left), H-H (center), and L-H (right). Error bars of the data points denote standard error. Solid lines show cumulative Gaussian functions fitted to the data. Dashed lines denote 25%, 50%, and 75% values. Horizontal error bars around the estimated 50% points denote 68% confidence intervals. RMSE values represent the root-mean-square error for evaluating the goodness of fit.

Stimuli were generated using Psychopy 2020.1.2 (RRID:SCR 006571, Peirce et al., 2019) based on Python 3.6. Each stimulus presentation consisted of a reference stimulus, a comparison stimulus, and a color gradient bar as a hue sequence reference (Fig. 2b). This color gradient was introduced to provide a direction of hue change, because, in contrast to orientation judgments, for hue there is no perceptual quality corresponding to clockwise vs. counterclockwise rotation. The gradient bar indicated from left to right a counterclockwise direction of hue changes along the color space azimuth (Fig. 2a-b). Both reference and comparison stimuli consisted of arrays of 16 circular patches with diameters of 0.75° of visual angle, evenly spaced on a 4 *⇥* 4 grid extending 3° *⇥* 3° of visual angle. Within each stimulus, the circular patches were randomly positioned on the grid of the array. The gradient color bar had an extent of 12° *⇥* 1.5° of visual angle.

On a neutral gray background (46.01° *⇥* 29.68°), a fixation cross (0.6° *⇥* 0.6°) was presented at the screen center. The gradient color bar was presented on the upper part of the screen, 5° above the center. Reference and comparison stimuli were placed 3° left and right from the fixation cross, with the sides of reference and comparison stimulus assigned in a pseudo-random fashion such that their positions were balanced within each session.

For both reference and comparison stimuli, the hues of the circular patches either were identical (low-noise stimuli) or were drawn from a uniform distribution with a range (noise level) that was individually pre-determined for each subject to yield thresholds twice as large as the low-noise stimuli (high-noise stimuli, see below). The reference hues, *θ_r_*, were evenly spaced along the azimuth of color space at 45° intervals from 22.5° to 337.5°. For each *θ_r_*, the corresponding gradient color bar was filled by 45 evenly spaced hue angles from *θ_r_* - 30° to *θ_r_* + 30°.

### 2.3 Procedures

During the experiment, the participant sat in a dimly lit room and viewed the display binocularly from a viewing distance of 57 cm. A central fixation cross was displayed and participants had to maintain their eyes to the fixation throughout the entire trial.

Each trial started with the presentation of the gradient color bar to indicate the range of hues to be tested. After 500 ms, the reference stimulus and the comparison stimulus were presented simultaneously for 500 ms, followed by a 500-ms full-screen checkerboard pattern of random chromatic squares (1.3° *×* 1.2°) to prevent afterimages of the stimuli. Participants were instructed to compare the two stimuli and indicate whether the ensemble hue averages of the arrays from left to right matched the direction of color change shown by the gradient color bar. Responses were given by pressing the up or down arrow key with the right hand. There was no time constraint on the response. Average response times were within one second.

For a given reference stimulus, its average hue and corresponding gradient color bar remained unchanged across trials, while the hue difference between the reference hue and the average hue of the comparison stimulus was adjusted by a 1-up/2-down staircase. Two same-noise conditions, low-noise versus low-noise (L-L) and high-noise versus high-noise (H-H), and one cross-noise condition, low-noise versus high-noise (L-H), were tested in the experiments. In the cross-noise condition, the reference stimulus was always the low-noise stimulus.

Each session consisted of 128 trials, in which the first 8 trials were warm-up trials with random hues and were excluded from further analysis. Within each session, two reference stimuli with 180° di↵erence in their mean hue angles, were randomly interleaved. Prior to the formal experiment, participants completed four practice sessions with feedback about the correctness of their responses given as text emojis (”:)” or ”:(” for correct and incorrect responses, respectively). Each participant performed 5760 trials divided into 24 conditions (8 reference stimuli x 3 comparison conditions) over 48 sessions.

As in the study by Girshick et al. (2011), prior to the main experiment, we determined each participant’s level of hue variance for the high-noise stimulus such that H-H discrimination thresholds were roughly 2 times larger than L-L discrimination thresholds. A reference hue angle of 135° was used because thresholds had intermediate values for this hue. For each participant, we measured the discrimination threshold (*d_L_*) of this stimulus in the L-L condition by fitting a cumulative Gaussian function to the psychophysical data. Next, we fixed the reference and comparison stimuli at 135° and 135°+2*d_L_*, respectively, and used a 1-up/2-down staircase to adjust the amount of the hue noise of both stimuli. The noise level was chosen that yielded 75% correct performance according to the psychometric function. The determined noise levels (21.2°, 20.1°, 24.5°, 20.1°, 22.2°, and 24.4° for 6 subjects, respectively) were used for the high-noise stimuli in the main experiment.

### 2.4 Data Analysis

#### 2.4.1 Estimation of perceptual variability and bias

The psychophysical data were analyzed separately for each subject. In addition, data for a hypothetical average observer were obtained by pooling all subjects’ data. For the data of each condition, we fitted a cumulative Gaussian function (Fig. 2c) using non-linear least square minimization with the Nelder-Mead algorithm (Gao and Han, 2012) and determined its mean, representing the point of subjective equality (PSE) and the standard deviation, representing the just noticeable difference (JND). PSE and JND thus reflected perceptual bias and perceptual variability, respectively.

#### 2.4.2 Estimation of the measurement distributions and the likelihood functions

The measurement distribution is the conditional distribution *p*(*m|θ*) and corresponds to the likelihood of a sensory measurement *m* given a particular stimulus hue angle *θ*. For each stimulus *θ* we estimated the measurement distribution as a von Mises distribution with a peak at *θ* and the concentration parameter *κ_θ_*, thus

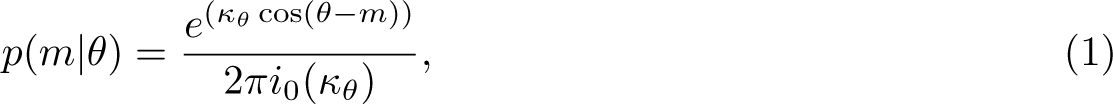

where *i*_0_(*κ_θ_*) is the modified Bessel function of order 0. The concentration parameter *κ_θ_* represented the measurement noise and was converted from the corresponding perceptual variability *J*(*θ*) by *κ_θ_* = *J^-2^*(*θ*). We estimated *J*(*θ*) by fitting each subject’s same-noise JNDs as a sine mixture function of the hue angle *θ*:

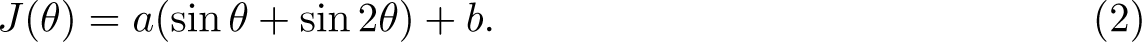

With periods of 180° and 360°, the two sine functions allow capturing both the periodicity and asymmetry of the JND patterns. The parameters *a* and *b* were estimated using non-linear least squares minimization.

To avoid any constraint of predefined shapes in estimating the likelihood, we adopted a sampling method using a pre-computed two-dimensional function (Girshick et al., 2011), where one dimension represented measurement distributions centered on particular stimulus hue angles *θ*, and the other dimension represented likelihood functions of *θ* given particular measurements *m*. Thus, a single measurement from the measurement distribution results in a likelihood corresponding to a horizontal slice of the two-dimensional function.

#### 2.4.3 Estimation of the priors

As the model of the prior *p*(*θ*) we chose a von Mises function (Eq. 1), which guaranteed that the prior had a period of 360° and an integral of 1. We determined the prior by fitting the estimation of a Bayesian observer to the behavioral data under the cross-noise condition (Girshick et al., 2011). We assume that the Bayesian observer encodes with sensory noise and gives distributed measurements *m*(*θ*) for repeated presentation of the same stimulus *θ*. Each measurement leads to a likelihood function, which is multiplied by the prior to obtain a maximum a posterior (MAP) estimate at the decoding phase. Note that, an alternative to the MAP estimate is the mean of the posterior; however, we opted for the MAP estimate in alignment with the approach outlined by Girshick et al.. The MAP estimate thus represents the observer’s estimate *θ*^^^ of the stimulus *θ*. Therefore, the measurement distribution of a stimulus results in a distribution of MAP estimates. The discrimination task was simulated by comparing two MAP estimate distributions according to signal detection theory (Green and Swets, 1966), yielding a single point on the simulated psychometric function.

For each participant, we simulated 1000 trials for each stimulus pair of the cross-noise comparison data. For every paired low-noise and high-noise stimuli, 1000 samples each were drawn from two corresponding measurement distributions with centers at *m_L_* and *m_H_*, and concentration parameters *κ_L_* and *κ_H_*, respectively. Each sample generated a likelihood function that was combined with the prior and led to an estimate of the stimulus. The two distributions formed by the 1000 estimates each were compared, resulting in a response probability given the corresponding stimulus pair. We then obtained a model-generated psychometric function by fitting a cumulative Gaussian function to the simulated data.

We evaluated the prior model by computing the likelihood of the cross-noise data given the model-generated psychometric function. The optimal parameters of the prior for each subject were estimated by maximizing the overall likelihood. We performed bootstrapping on each subject’s binary responses for each stimulus pair 100 times and estimated the priors given the bootstrapped data. The standard deviation of the 100 estimated priors was taken as the confidence interval of the optimized prior. We further assessed the model by comparing its performance with the performance of a model with a uniform prior. We evaluated the performance of the model by a normalized difference of log-likelihood to the model with the uniform prior,

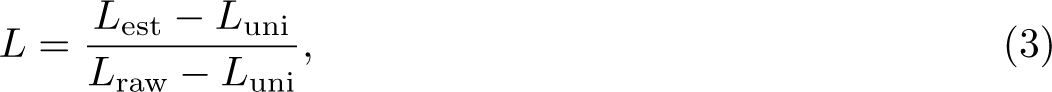

where *L*_est_ and *L*_uni_ represent the log likelihoods of the models with the estimated prior and the uniform prior, respectively, and *L*_raw_ represents the log likelihoods of the raw psychometric fits. Thus, *L* = 0 corresponds to the model with a uniform prior and *L* = 1 corresponds to the raw psychometric fits. Given the difference in the degrees of freedom between the estimated prior and the uniform prior, we also calculated the Akaike Information Criterion (AIC) scores (Akaike, 1998) of the models.

To validate the unimodality of the prior model, we also modeled two alternative priors by normalized sine functions with periods of 360° and 180°, respectively, and determined the prediction performance of the observer models with these priors.

## 3 Results

### 3.1 Behavioral Results

We used two-alternative forced-choice discrimination experiments to measure perceptual variability and bias in hue perception. We tested three conditions for each measured hue: two same-noise conditions (low-noise versus low-noise, L-L, and high-noise versus high-noise, H-H), and one cross-noise condition (low-noise versus high-noise, L-H). The same-noise conditions enabled measuring the perceptual variability of stimuli with specific noise levels, while the data in the cross-noise condition could potentially reveal the effect of a prior through a cross-noise bias (Fig. 1).

We fitted the psychometric data for hue discrimination across varying noise conditions with cumulative Gaussian functions (see Fig. 2c for examples). The goodness of the psychometric fit was measured by the root-mean-square errors (RMSE) and was comparable across subjects (0.05 *±* 0.02 for the L-L condition, 0.08 *±* 0.02 for the H-H condition, and 0.07 *±* 0.02 for the L-H condition). Based on the estimated psychometric function, we used the standard deviation of the cumulative Gaussian function as the just noticeable difference (JND) and calculated the point of subjective equality (PSE) at the threshold of 50%. These values reflect the discrimination thresholds and bias of each participant, respectively. In addition, we determined the results for a hypothetical average subject by pooling the data from all subjects. This average subject showed an average performance: there was no significant difference between the average subject’s psychophysical estimates and the mean values of all subjects’ psychophysical measurements (sign test *p* = 0.44 for discrimination thresholds, sign test *p* = 0.68 for biases).

#### 3.1.1 Discrimination thresholds

Under the same-noise conditions, discrimination thresholds as a function of hue angle typically exhibited a bimodal pattern (Fig. 3a). On average, across subjects, the L-L condition had two local minima at hue angles of 101.3° *±* 22.5° and 298.1° *±* 28.8° (see Fig. S1 and S2), and two local maxima occurred at 49.5° *±* 24.6° and 170.4° *±* 21.9°. Fitting a sine mixture model to the discrimination thresholds yielded similar hue angles corresponding to these extrema (local minima at 97.3° *±* 28.4° and 290.7° *±* 18.1°, and local maxima at 39.4° *±* 47.1° and 186.7° *±* 23.6°, averaged across subjects; Figure 4a-b, also see Fig. S4a).

**Figure 3:**
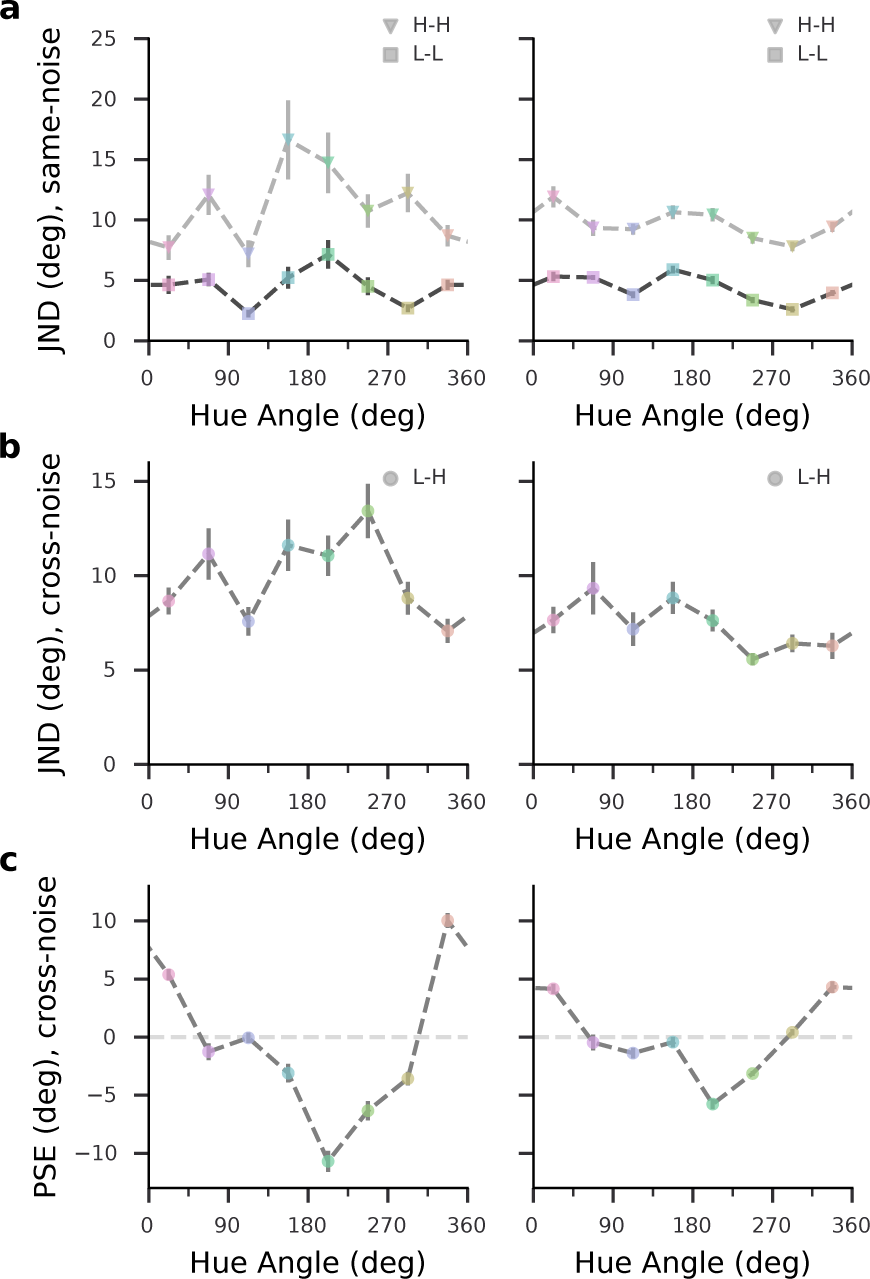
Experimental data for subject S3 (left) and the average subject (right). (a) Hue discrimination thresholds (JNDs) under the same-noise condition. (b) Hue discrimination thresholds under the cross-noise condition. (c) Biases under the cross-noise condition, measured as hue angle di↵erences between the high-noise and low-noise stimuli at the PSE. Bars denote one standard error of the estimates.

**Figure 4:**
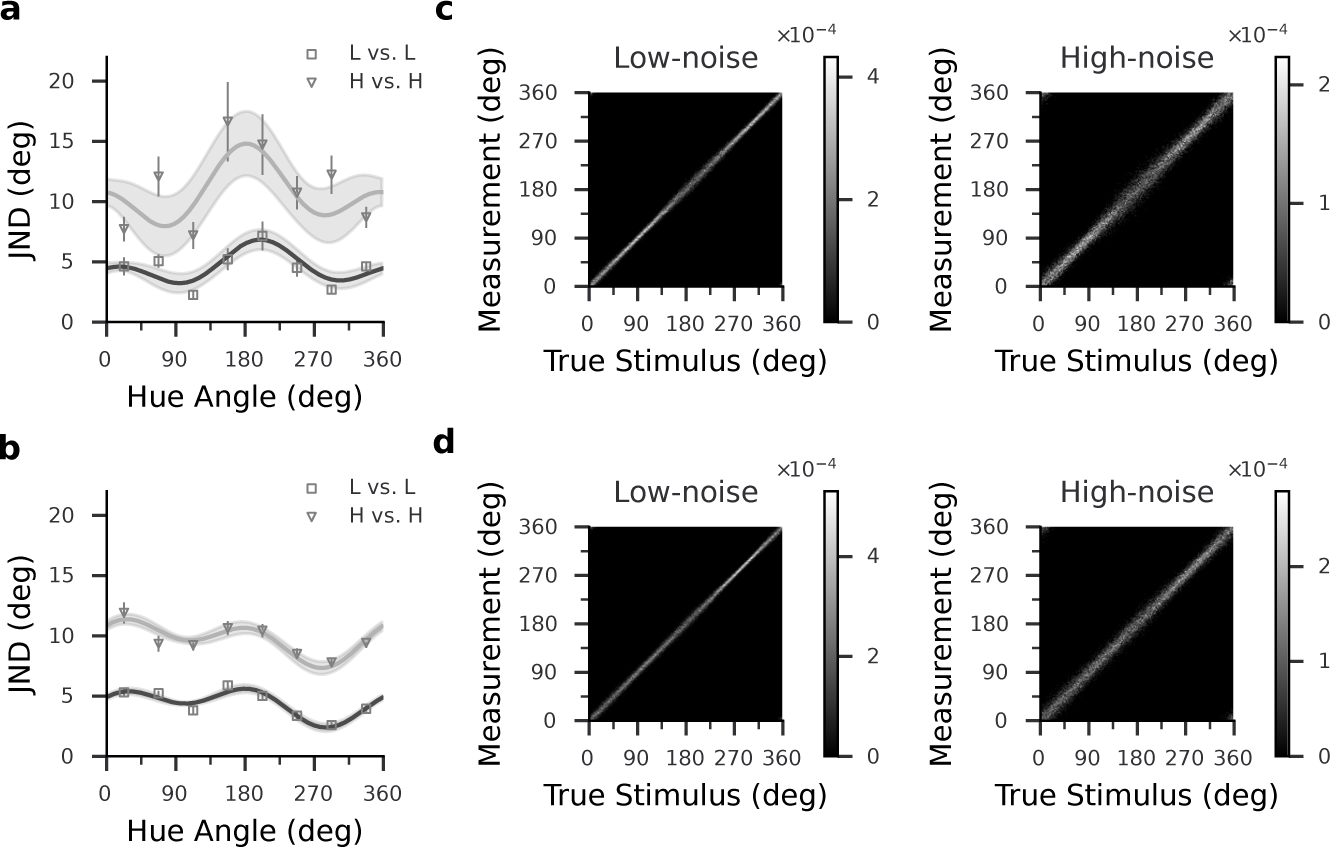
Threshold fits and estimated likelihood functions in the same-noise conditions. (a-b) Fitted JNDs of subject S3 (a) and the average subject (b). JND estimates are from the data shown in Fig. 3, error bars indicate one standard error. The dark and light gray lines are the fitted JNDs for the L-L and the H-H conditions, respectively. The gray shaded area indicates 68% confidence intervals of fitted JNDs. (c-d) Estimated likelihood functions of subject S3 (c) and the average subject (d). Each horizontal slice of the two-dimensional function represents a likelihood function of stimulus hue angle given a particular measurement *m*, and each vertical slice represents a measurement distribution centered on a particular *θ*. The gray level represents corresponding probability densities.

The bimodal pattern of discrimination thresholds, with lowest thresholds near an oblique axis connecting blue and yellow, is in line with results of previous studies (Danilova and Mollon, 2010; Witzel and Gegenfurtner, 2013). In the H-H condition, thresholds were significantly higher than in the L-L condition (sign test *p< .*001) but the bimodal pattern persisted, however two of the subjects (S2 and S6) showed inconsistent maxima and minima between the conditions (Fig. S1). The cross-noise condition (L-H) typically yielded intermediate discrimination thresholds (Fig. 3b). Of the L-H discrimination thresholds, 82.5% were higher (sign test *p< .*001) than the corresponding L-L discrimination thresholds, and 90.0% were lower (sign test *p < .*001) than the H-H discrimination thresholds.

#### 3.1.2 Biases

We observed non-negligible bias only under the cross-noise condition (L-H) for all subjects (Fig. 3c; also see Fig. S3 for the same-noise bias results), which matches our prediction based on the Bayesian inference framework: assuming the same prior for perceptual inference, biases would only di↵er among stimuli with di↵erent noise levels. Given that in the cross-noise condition the low-noise stimulus always served as the reference stimulus, with fixed hue across trials, the cross-noise bias shown in Fig. 3c represents the perceptual bias of the high-noise stimulus relative to the low-noise stimulus. Averaged across subjects, the biases showed a minimum of *-*5.6° *±* 3.1° and a maximum of 5.5° *±* 2.9°. Among the biases, 88.9% of the negative values corresponded to hue angles within the range of 112.5° to 292.5°, and 85.7% of the positive values were at the hue angles smaller than or equal to 112.5°, or larger than or equal to 292.5°. Subjects’ biases showed zero-crossings at the hue angles within 45° around 112.5° and within 45° around 292.5°, except for one subject whose zero-crossings did not fall within the hue angles of 112.5° *±* 45°. Around the zero-crossings, biases were attractive towards blue hues (near 112.5°) and repulsive away from yellow hues (near 292.5°). Most subjects exhibited two zero-crossings, around 112.5° and 292.5°, respectively. The two subjects that had inconsistent discrimination thresholds across conditions showed additional bias zero-crossings around 157.5° (see S2 and S6 in Fig. S1). According to the Bayesian observer model, perceptual bias arises when the prior is non-uniform over the stimulus space and misaligned with sensory measurements. This prediction matches our observations of non-zero cross-noise bias, suggesting non-uniformity of the perceptual prior.

Since our color space is not perceptually uniform, we expect some bias to arise from the variation of discrimination thresholds. If changes in thresholds occur over a scale of hue angles comparable to the hue range of the noisy stimulus arrays, then the perceived ensemble average for a given stimulus may vary depending on the noise level. To determine the contribution of threshold non-uniformities to observed bias, we simulated the effect of non-uniform thresholds when the paired cross-noise stimuli had identical central hues of the ensemble. For each subject, we multiplied the hue angle differences between the hues in the high-noise stimuli and the central hue of the ensemble by the inverse of the fitted JNDs (see Modeling Results and Fig. 4a-b). Thus, we mapped the stimuli to a scale where hue differences were represented as multiples of discrimination thresholds and then computed the hue averages of the scaled stimuli. This resulted in biases that varied systematically with hue angle, corresponding to the variation in discrimination thresholds. However, their magnitudes were considerably smaller than the observed cross-noise bias (Fig. S1). Averaged across observers, the biases arising from threshold non-uniformities showed maximal magnitudes of 0.91° *±* 0.78°, that is, less than 20% of the magnitudes of the experimentally measured biases. Thus, the variation of discrimination thresholds had a negligible influence on the cross-noise biases.

Taken together, we found that hue discrimination thresholds followed a bimodal pattern, with observers showing the best discrimination for bluish and yellowish stimuli. Introduction of chromatic noise resulted in increases in discrimination thresholds and cross-noise biases. The observed biases were attractive towards blue and repulsive from yellow, which is indicative of non-uniform priors.

### 3.2 Modeling Results

To determine priors employed by observers in hue judgments, we used a Bayesian ideal observer model and optimized the parameters of its prior to account for the data from our human observers. The model connects two behavioral measurements (discrimination threshold and bias) to two Bayesian components (likelihood and prior). Specifically, the stimulus uncertainty was propagated from the measurement distribution to the posterior distribution, resulting in perceptual variability (Girshick et al., 2011). In line with this model, measurement distributions and likelihood functions were computed from the fitted same-noise variabilities. When stimulus noise increased the threshold, the widths of the corresponding measurement distribution and likelihood function were also increased (Fig. 4, see also Fig. S4). We assumed the ideal observer’s estimates of a particular stimulus *θ* corresponded to maximum a posteriori (MAP) estimates *θ*^^^ resulting from multiplying the likelihood function with a prior at the Bayesian decoding stage. We simulated each subject’s cross-noise data by comparing each pair of MAP estimate distributions (*θ*^^^*_L_* and *θ*^^^*_H_*). The prior was modeled as a von Mises function, and the prior parameters were obtained by maximizing the likelihood of the experimental data given the simulation-generated psychometric function. Subjects’ priors peaked in the second quadrant, with the exception of the two subjects with inconsistent bias patterns, whose priors were shallow and estimates noisy (Fig. S5). For the average data, the estimated prior showed the highest values at 115.2° *±* 19.9° (Fig. 5). The cross-subject average of the hue angle where the prior peaked was 107.3° *±* 26.7° (SE), that is, close to blue along the axis of variation of natural daylights (Webster et al., 2000; Mollon, 2006), which has a hue angle of 112° in our color space.

**Figure 5:**
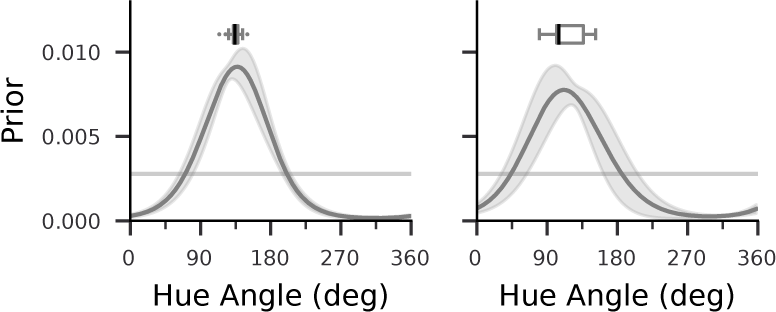
Estimated prior for subject S3 (left) and the average subject (right). The gray shaded area indicates *±*1 one standard deviation of 100 bootstrapped estimates. The boxes above the curves indicate the first quartile to the third quartile of the peak locations of the 100 bootstrapped estimates, with the black line at the median. The whiskers extend from the box by 1.5*×* the interquartile range (IQR). Flier points indicate values beyond the range of the whiskers. The light gray horizontal line represents the uniform prior.

To assess the e↵ectiveness of the encoding-decoding model with the estimated prior, we compared its performance in predicting the cross-noise bias against a model with a uniform prior. The model with the estimated prior was found to be better at predicting both the sign and amplitude of the cross-noise bias than the model with the uniform prior (Figure 6a-b). Across all subjects, the normalized log-likelihoods indicated that the estimated prior outperformed the uniform prior (Fig. 6c). Furthermore, when comparing the two prior models using Akaike Information Criterion (AIC) scores (Akaike, 1998), we found that the estimated prior consistently performed better than the uniform prior (Table S1). Additionally, modeling the prior as a normalized sine function yielded similar results, with the estimated prior peaking around the blue hue (119.1° *±* 28.6° (SE), averaged across subjects, see Fig. S7). Note that the sine function had a constrained period of 360° and thus exhibited a single peak over the hues. Its prediction performance was better than a bimodal sine function with period 180° (Fig. S7), which confirmed the unimodality of the prior.

**Figure 6:**
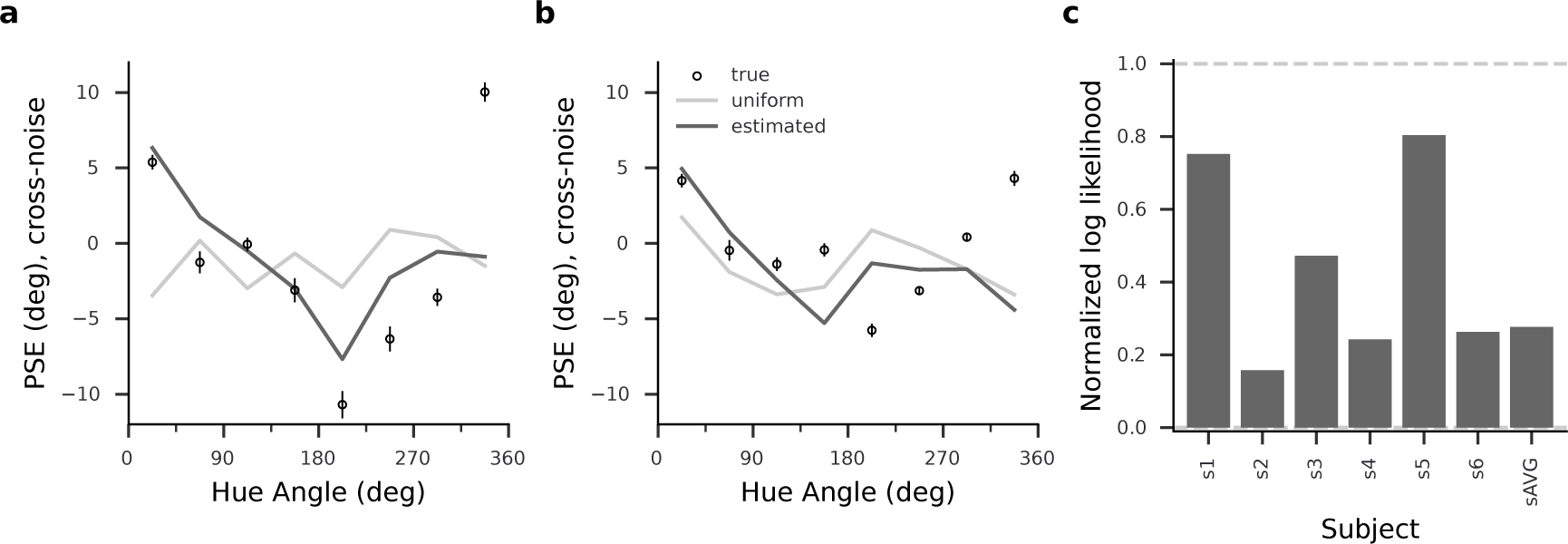
Comparison of priors. (a-b) Cross-noise biases with predictions from estimated prior (black lines) and uniform prior (gray lines) for subject S3 (a) and the average subject (b). Circles represent the cross-noise biases shown in Fig. 3. (c) Normalized log-likelihoods of predictions using the estimated prior for all subjects, including the average subject (sAVG). Values greater than 1 indicate prediction better than raw psychometric fits, and values greater than 0 indicate prediction better than the model with uniform priors.

In summary, these results indicate that the observers used a non-uniform prior related to the blue-yellow axis. Specifically, the prior showed the highest probability at blue and the lowest probability at yellow.

## 4 Discussion

Our study aimed to investigate the perceptual characteristics of hue discrimination and to identify an internal prior that contributes to hue perception. The experimental results showed that the lowest discrimination thresholds and smallest biases occurred for stimuli near a perceptual blue-yellow hue axis. In addition, cross-noise perceptual biases were attractive towards blue and repulsive from yellow, which was explained by a Bayesian observer model with a prior that favored blue. Our study extends the Bayesian perspective on perception to the domain of color and provides evidence of a systematic bias in color perception related to natural daylights. In contrast to previous attempts, where it had turned out difficult to determine adequate priors to explain human performance in color judgments (Brainard et al., 2006), our study presents an approach that recovers a prior that explains hue perceptual bias and reflects natural color statistics.

### 4.1 Ensemble hue perception

Our psychophysical results showed that subjects could effectively integrate the information over noisy hue ensembles, which agrees with previous findings on ensemble hue perception (Maule et al., 2014; Webster et al., 2014; Maule and Franklin, 2015; Virtanen et al., 2020). Specifically, our data confirm that hue averaging does not require a spatial configuration with abutting hue elements (Virtanen et al., 2020).

Previous studies have indicated that categorical effects may influence the percept of hue ensembles (Maule et al., 2014). However, discrimination thresholds in our experiments were approximately equal between yellow and blue (*p* = .35), despite their categories spanning different hue angle ranges (Webster et al., 2000; Hansen et al., 2007; Witzel and Gegenfurtner, 2013). Moreover, observers were instructed to discriminate hue using an external physical reference instead of internal criteria, minimizing potential categorical effects from subjective color naming. Nevertheless, to identify any relationship of the priors with individual specifics in color vision, we compared the locations of priors and unique hue percepts of the individual subjects. We determined the unique hue locations in five subjects (see Supplementary Methods) and found the standard deviation in unique hue locations (5.31° *±* 2.05°, averaged across four unique hues) was much smaller than that in the prior peak locations (70.46°). For one observer (S1) whose blue and yellow unique hue locations deviated from other subjects’ settings towards larger hue angles, the corresponding prior peaked at a hue angle close to the average peak location of all subjects’ priors. An observer (S6) with a prior peaking at the largest hue angle among subjects did not show pronounced deviations from the other subjects in unique hue categories. Overall we did not find a correlation between the prior peak location and any of the unique hue locations (Pearson correlation *r* = *-*0.33*,p* = 0.59 for unique blue, *r* = *-*0.77*,p* = 0.12 for unique yellow, *r* = *-*0.09*,p* = 0.88 for unique red, *r* = *-*0.42*,p* = 0.47 for unique green). Thus, it is unlikely that the biases we observed could be attributed to categorical effects.

### 4.2 Relation between perceptual variability and bias

The two primary psychophysical measures - perceptual variability and bias - covaried in our experiments: both discrimination thresholds and biases were lowest near the blue-yellow axis and largest orthogonal to the blue-yellow axis. This finding is consistent with previous studies on orientation perception, which have shown a similar relation between threshold and bias minima occurring at cardinal orientations (Tomassini et al., 2010; Girshick et al., 2011).

However, as previous studies have reported, orientation stimuli with high discrimination thresholds or high variability, such as oblique orientations, can also be perceived with minimal bias (Tomassini et al., 2010; Girshick et al., 2011). Wei and Stocker (2017) presented a mathematical description of the relation between variability and bias, suggesting a proportional relationship between bias and the derivative of the square of the discrimination threshold, and found that the relation holds for many visual features, including orientation, motion direction, magnitude, and spatial frequency (Wei and Stocker, 2017). The relationship predicts that bias is minimal at the extrema of discrimination thresholds, with attraction towards the maxima and repulsion bias away from the minima of discrimination thresholds. According to this prediction, one would expect that the bias of perceived hues in our study would show four zero-crossings, corresponding to the number of maxima and minima in the bimodal pattern of discrimination thresholds. Specifically, the hue percept should be biased away from the blue-yellow axis, where discrimination thresholds are minimal, and attracted towards colors with high discrimination thresholds, such as reddish and greenish hues (Fig. S8). However, the measured cross-noise biases showed repulsion from yellow and attraction towards blue.

Potentially, our measurements might have missed to consistently identify some zero crossings, particularly around 157.5° where inter-subject variability occurred in biases (Fig. 3c) and some subjects showed more than two zero-crossings (Fig. S1). Even if this were the case, the attraction bias around blue in our results is nevertheless inconsistent with the prediction of the Wei-Stocker relation, which would predict that the bias should be repulsive away from blue. An alternative possibility that could explain the deviation of our results from the relation is a strongly skewed likelihood function. Intuitively, one would expect the bias to be zero when the prior of a Bayesian observer is uniformly distributed across the entire range of stimuli. However, the model with a uniform prior predicted non-negligible bias for some of our subjects (Fig. 6, see also S2 and S6 in Fig. S6). These results are likely related to the asymmetry of the likelihood function in the observer model (Wei and Stocker, 2015), which resulted from estimating the likelihood directly from the experimental data by sampling from a measurement distribution, a method also employed by Girshick et al. (2011). While it might be feasible to simultaneously model the likelihood and prior (Stocker and Simoncelli, 2006), we relied on the measurement distribution to ensure reliable likelihoods that captured the perceptual variability for stimuli with specific noise levels. A heavy-tailed likelihood might lead to a deviation of the posterior from the true stimulus, such that both likelihood shape and prior could contribute to perceptual bias (Stocker and Simoncelli, 2006; Wei and Stocker, 2015; Prat-Carrabin and Woodford, 2021). However, the asymmetric likelihood with a uniform prior generated less accurate predictions than the model with the non-uniform prior (Fig. S6). Thus, the key factor in yielding the systematic bias in our experiments was the non-uniformity of the prior.

Could the marked deviation from the Wei-Stocker relation indicate that the perception of color is governed by fundamentally different principles than other visual features? Wei and Stocker derived their relation under assumptions of efficient coding, specifically, that stimulus encoding maximizes the mutual information between stimuli and sensory representations. This is reasonable when considering visual features with simple stimulus correspondence, such as spatial features including orientation or motion direction. But given the multiple transformations of color signals in the visual system, perceptual judgments in the chromatic domain may be subject to more complex constraints. Sensory signals for color vision are encoded first in a cone-opponent fashion by the retinal circuitry. The resulting representation is the basis for the color space that is commonly used (MacLeod and Boynton, 1979; Derrington et al., 1984), including in our study. Retinal cone opponency decorrelates the photoreceptor signals and thus reduces redundancy (Ruderman et al., 1998; Lee et al., 2002).

However, it does not capture the distribution of natural chromatic signals to achieve maximal information (Wachtler et al., 2001; Kellner and Wachtler, 2013). Specifically, retinal cone opponency does not align with the variation of natural illumination, as is reflected by the non-cardinal orientation of the daylight axis in cone-opponent color space (Mollon, 2006). A sensory representation better matched to the distribution of natural chromatic signals appears only at a later stage, by the transformation of color signals in the visual cortex, where the precortically separated cone-opponent signals (Chatterjee and Callaway, 2003) are mixed and a distributed code is achieved (Lennie et al., 1990; Wachtler et al., 2003; Kuriki et al., 2015; Li et al., 2022) that captures the oblique axis of variation of natural daylights (Wachtler et al., 2003; Lafer-Sousa et al., 2012). While color appearance judgments are likely based on the cortical representation, perceptual variability must be assumed to be influenced considerably by the signal-to-noise ratios at precortical stages (Vorobyev and Osorio, 1998). These different influences may result in thresholds and biases that appear inconsistent with the Wei-Stocker relation.

### 4.3 Hue perceptual non-uniformity and blue-yellow asymmetries

Perceptual quantities as measured in our study are not uniform when represented in a color space based on cone-opponency. Specifically, discriminability and variability vary in this color space (Boynton et al., 1986; Krauskopf and Gegenfurtner, 1992; Danilova and Mollon, 2010; Bosten et al., 2015; Klauke and Wachtler, 2015). Our results verified that such non-uniform discriminability also exists for hue ensembles with chromatic noise. Moreover, we found that biases in hue judgments varied with hue, showing minima near blue and yellow, which further reflected the non-uniformity. Notably, the non-uniformity arises not along the cardinal axes of precortical cone opponency, but with respect to the oblique axis connecting perceptual unique blue and yellow that coincides with the variation of natural daylight (Mollon, 2006). This axis has been found special for color vision in many respects, from the distribution of natural chromatic signals (Webster and Mollon, 1997; Wachtler et al., 2001; Webster, 2020) to neural processing (Wachtler et al., 2003; Lafer-Sousa et al., 2012) and perception including discrimination (Danilova and Mollon, 2010; Bosten et al., 2015), color induction (Klauke and Wachtler, 2015) and color constancy (Delahunt and Brainard, 2004; Pearce et al., 2014; Gegenfurtner et al., 2015; Lafer-Sousa et al., 2015; Weiss et al., 2017).

While blue and yellow appear symmetrical, in terms of their similarities in perceptual quantities and the coincidence with the daylight locus (Mollon, 2006; Webster, 2020), the bias in our results was attractive towards blue and repulsive from yellow, confirming asymmetries between blue and yellow (Webster, 2020). A previous study has demonstrated a systematic deviation towards blue when subjects adjusted yoked blue-green hue pairs to achieve an equal perceived mixture of binary hues (Webster et al., 2014). The deviation occurred for unique blue settings and not for other unique hues, which matches the unimodality of the systematic prior revealed in our results. Moreover, the deviation only occurred in the blue-green settings, while no conspicuous bias arose in the mixture consisting of yellow hues, which strongly evidenced asymmetries between blue and yellow.

A special role of blue has been observed previously in color constancy: bluish illumination results in higher color constancy than other chromatic illuminations (Delahunt and Brainard, 2004; Pearce et al., 2014; Winkler et al., 2015; Radonjic et al., 2016; Weiss et al., 2017; Aston et al., 2019). An explanation for this so-called “blue bias” was that the illumination sensitivity threshold is higher for blue than for other colors (Pearce et al., 2014; Radonjic et al., 2016), which would be in line with the hypothesis that the visual system may adapt to the natural environment and be least sensitive to the illumination changes that are most likely to occur (Aston et al., 2019). Alternatively, given the color distribution of lighting and shadows in natural scenes, the blue bias may emerge from the observer’s tendency to infer blue tints as illuminants (Winkler et al., 2015). Although these explanations do not reconcile, they commonly imply an environmental account of the blue-yellow asymmetry: color vision may be adapted to natural spectra and expect blue illumination as a dominant feature in natural conditions.

In line with these interpretations, our observer model attributes the blue-yellow asymmetries at the behavioral level to a unimodal prior that peaks at blue. Notably, the unimodal prior outperforms bimodal or uniform priors in predicting the perceptual biases (Fig. S7). Our model is in line with the notion that perception, and in particular color perception, is shaped by the regularities of the sensory environment (Shepard, 1992), but also suggests an asymmetry in natural daylight. As Mollon (2006) has pointed out, the clear sky appears unique blue, which strongly indicates that the light of the sky, resulting from the fundamental physical process of Rayleigh scattering, might provide a stable reference to which color perception is anchored. Thus, in keeping with other Bayesian approaches, our results suggest that human perception internalizes the natural sensory statistics and incorporates prior knowledge into the processing of sensory information.

## Author Contributions

**Yannan Su**: Methodology, Software, Formal analysis, Investigation, Data Curation, Visualization, Writing - Original Draft, Writing - Review & Editing. **Zhuanghua Shi**: Writing - Review & Editing, Supervision. **Thomas Wachtler**: Conceptualization, Methodology, Writing - Review & Editing, Supervision.

## Acknowledgments

Supported by DFG (RTG 2175 ”Perception in Context and its Neural Basis”) and Bernstein Center for Computational Neuroscience Munich. We thank all participants for participating in the experiments.

