## Supplementary Materials for "A Bayesian observer model reveals a prior for natural daylights in hue perception"

### Data Availability

The data and code can be found at G-Node: https://doi.org/10.12751/g-node.rp2ft3.

### Supplementary Methods

#### Unique Hue Measurements

We measured unique hue locations to investigate whether the priors were related to the participants' color categories. Five of the recruited participants participated in this supplementary experiment. The stimulus was a 10° × 10° colored rectangle presented on a neutral gray background. Participants were instructed to adjust the hue of the stimulus by moving the mouse until the color appeared a unique hue. For example, in a trial measuring subjective unique blue, the task was to make the color appear "blue and neither reddish nor greenish". Participants adjusted the stimulus along the azimuth of the color space. There was no time constraint on the response. Participants confirmed the adjustment and terminated the trial by pressing the space bar on the keyboard. Four unique hues (red, blue, green, and yellow) were tested with 30 repeated trials each. The initial hue angle of the stimulus color was randomly chosen from a range of 40° around a guessed value of the unique color (0°, 110°, 220°, and 315° for red, blue, green, and yellow, respectively) based on previous studies (Webster et al., 2000). To prevent after-image and adaption, a full-screen 500-ms checkerboard pattern of chromatic squares was presented after the subject's response, and all trials were randomly interleaved.

### Supplementary Figures and Tables

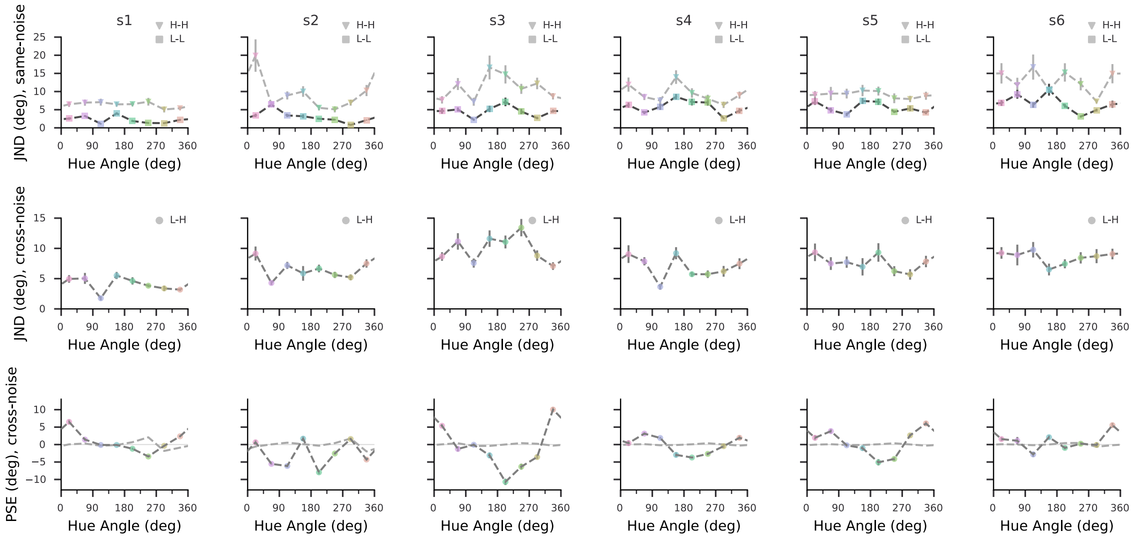


Figure S1: Experimental data for each subject (each column, respectively). Top: hue discrimination thresholds (JNDs) under the same-noise condition. Middle: hue discrimination thresholds under the cross-noise condition. Bottom: biases under the cross-noise condition, measured as hue angle differences between the high-noise and low-noise stimuli at the PSE. Bars denote one standard error of the estimates. Light gray dashed lines represent the bias induced by local non-uniformity of discrimination (see Results).

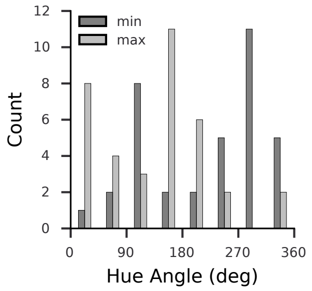


Figure S2: Distribution of stimulus hue angles corresponding to the two local minima (dark gray bars) and two local maxima (light gray bars) of the discrimination thresholds. Data are from all subjects and all conditions.

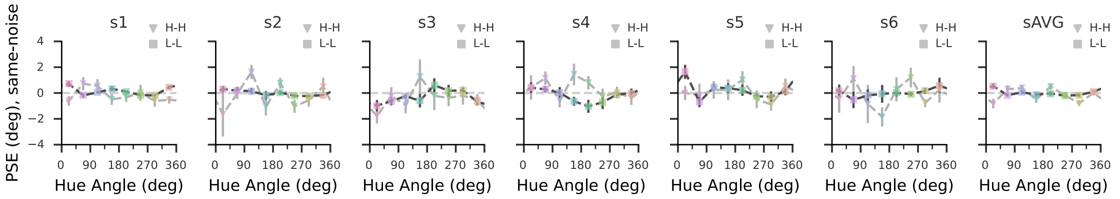


Figure S3: Same-noise bias for each subject (each column, respectively). Bars denote one standard error of the estimates.

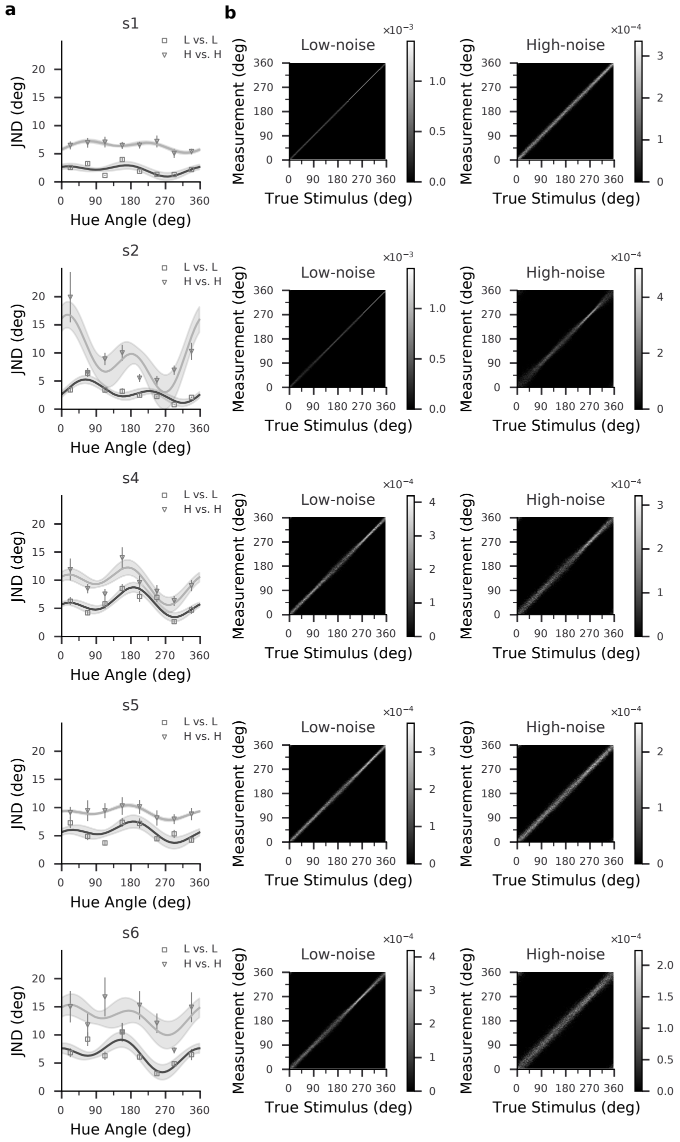


Figure S4: Threshold fits and estimated likelihood functions in the same-noise conditions for subjects S1, S2, S4, S5, and S6 (five rows, respectively). (a) Fitted JNDs of the subjects. JND estimates are from the data shown in Fig. S1, error bars indicate one standard error. The dark and light gray lines are the fitted JNDs for the L-L and the H-H conditions, respectively. The gray shaded area indicates 68% confidence intervals of fitted JNDs. (b) Estimated likelihood functions of the subjects. Each horizontal slice of the two-dimensional function represents a likelihood function of stimulus hue angle $\theta$ given a particular measurement $m$, and each vertical slice represents a measurement distribution centered on a particular $\theta$. The gray level represents corresponding probability densities in the L-L condition(”Low-noise”, left) and in the H-H condition (”High-noise”, right).

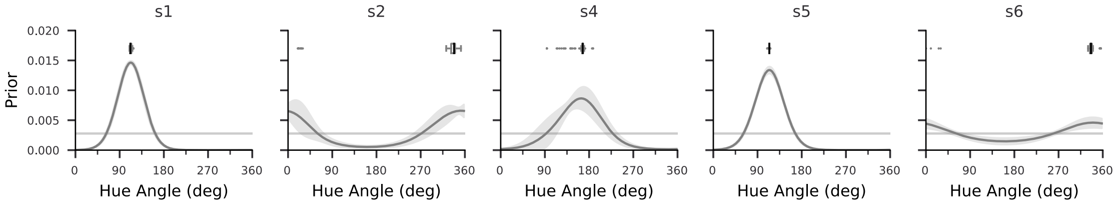


Figure S5: Estimated priors for subjects S1, S2, S4, S5, and S6 (five columns, respectively). The gray shaded area indicates ±1 the standard deviation of 100 bootstrapped estimates. The boxes above the curves indicate the first quartile to the third quartile of the peak locations of the 100 bootstrapped estimates, with the black line at the median. The whiskers extend from the box by 1.5× the interquartile range (IQR). Flier points indicate values beyond the range of the whiskers. The light gray horizontal line represents the uniform prior.

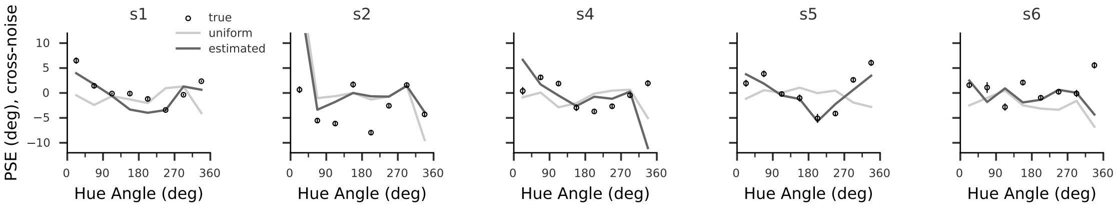


Figure S6: Cross-noise biases with predictions from estimated priors (black lines) and uniform prior (gray lines) for subjects S1, S2, S4, S5, and S6 (5 columns, respectively). Circles represent the cross-noise biases shown in Fig. S1.

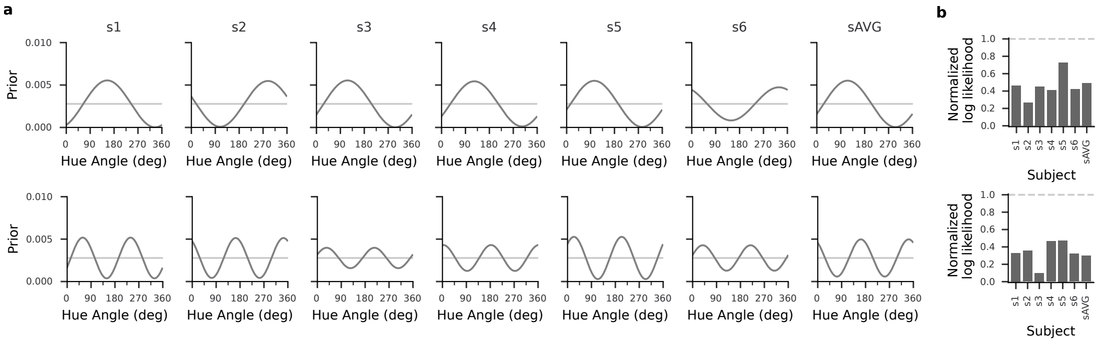


Figure S7: Alternative prior models. Estimated priors as sine functions for all subjects (including the average subject, sAVG). The periods of the estimated sine-shape priors were 360° (top) and 180° (bottom), respectively. (a) Estimated sine-shaped priors for all subjects (7 columns, respectively). The light gray line represents the uniform prior. (b) Normalized log-likelihoods of predictions using the estimated sine-shape prior for all subjects.

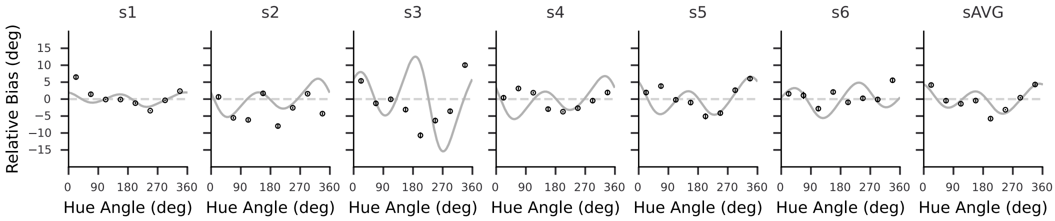


Figure S8: Cross-noise bias predicted according to the Wei and Stocker (2017) relation. Bias curves (gray) were calculated using the mathematical relation between perceptual variability and bias described by Wei and Stocker (2017). Black dots represent the measured cross-noise biases (same data as in Fig. 3 and Fig. S1).

| Subject | Estimated Prior | Uniform Prior |
|---|---|---|
| s1 | 2,097 | 2,724 |
| s2 | 2,473 | 2,560 |
| s3 | 2,365 | 2,633 |
| s4 | 2,311 | 2,388 |
| s5 | 2,421 | 2,988 |
| s6 | 2,318 | 2,379 |
| sAVG | 2,458 | 2,539 |

Table S1: Akaike Information Criterion (AIC) scores of the estimated and uniform priors for all subjects.
